# Fluorescent substances in cotton fibers treated with a labeling probe

**DOI:** 10.1101/2024.05.13.593715

**Authors:** Shigeru Tamogami, Akio Watanabe, Ganesh K. Agrawal, Randeep Rakwal

## Abstract

Usability of the labeling probe, previously developed to uncover the targets of cellulose synthase (CESA) inhibition herbicides, was examined with fibers and cultured ovules of the cotton plant (*Gossypium hirsutum*). Using this probe, various fluorescent substances were found with attributes, such as bands, particles and other unique structures. Elucidation of these fluorescent substances might provide hints to understand the mechanism of cellulose biosynthesis in the cotton fibers and ovules, as well as to distinguish features of the cotton fibers.

## Introduction

Cellulose is one of the most useful and abundant natural polymers on the planet. For plants, cellulose is an important component of the cell walls that support the structure of the cells. It has been widely known that cellulose biosynthesis is essential for the cell elongation, growth and division. Plants biosynthesize cellulose from glucose through the condensation reaction, which is catalyzed by enzymes, termed as cellulose synthases (Anderson C and Kieber J, 2020). To be noted, some inhibitors, which are suggested to target cellulose synthases, have been utilized as herbicides (Larson R and McFarlane H, 2021). We have designed and synthesized a labeling probe based on such herbicides (Tamogami et al., 2021). We tested usability of this probe and found that fluorescent substances were localized along the cell walls of *Arabidopsis* root epidermal cells, where cellulose biosynthesis would be active (Fig **1** [**A**], unpublished result).

To ensure the usability, we applied this probe to cotton plants, which produce a large amount of cellulose in the cotton fibers. As a result, we determined that characteristic fluorescent substances were localized in the cotton fibers treated with the probe. Moreover, we found unique fluorescent structures, in the cultured ovules treated with the probe. We hope that our present results would contribute to cotton science, and elucidation of these fluorescent substances are now under way.

## Methods

### 1. Materials and reagents

Cotton plants (*Gossypium arboreum* seeds purchased from TOHOKU SEED CO., LTD., Utsunomiya, Japan) were grown from seeds during May to August, 2023 at the University field and in the green house during October, 2023 to May, 2024 (Akita, Japan). Flowers and cotton bolls of *Gossypium hirsutum* were gifts from the University experimental field center (seeds were purchased from MEDETASHI-SYUBYOU, Chikusei, Ibaraki Japan). Representative plant materials of the cotton boll and fibers (*G. hirsutum*) were shown in **Fig. 1** [**B**] and [**C**]. Cotton ovule cultures were prepared in a culture medium according to a protocol reported by (Tan et al., 2013). Fresh flowers were harvested, and 0 DPA (day post-anthesis) ovules were used to prepare the ovule cultures. Independent preparation of the cotton ovule cultures was performed at least 3 times and used for observations. Representative ovule cultures were shown in Fig. **1** [**D**]. Reagents to prepare the culture medium were purchased from Sigma-Aldrich Japan. A special grade of dimethylsulfoxide (DMSO, FUJIFILM Wako Pure Chemical Corporation, Japan) was used in the preparation of the probe reagent solution.

**Fig. 1.**
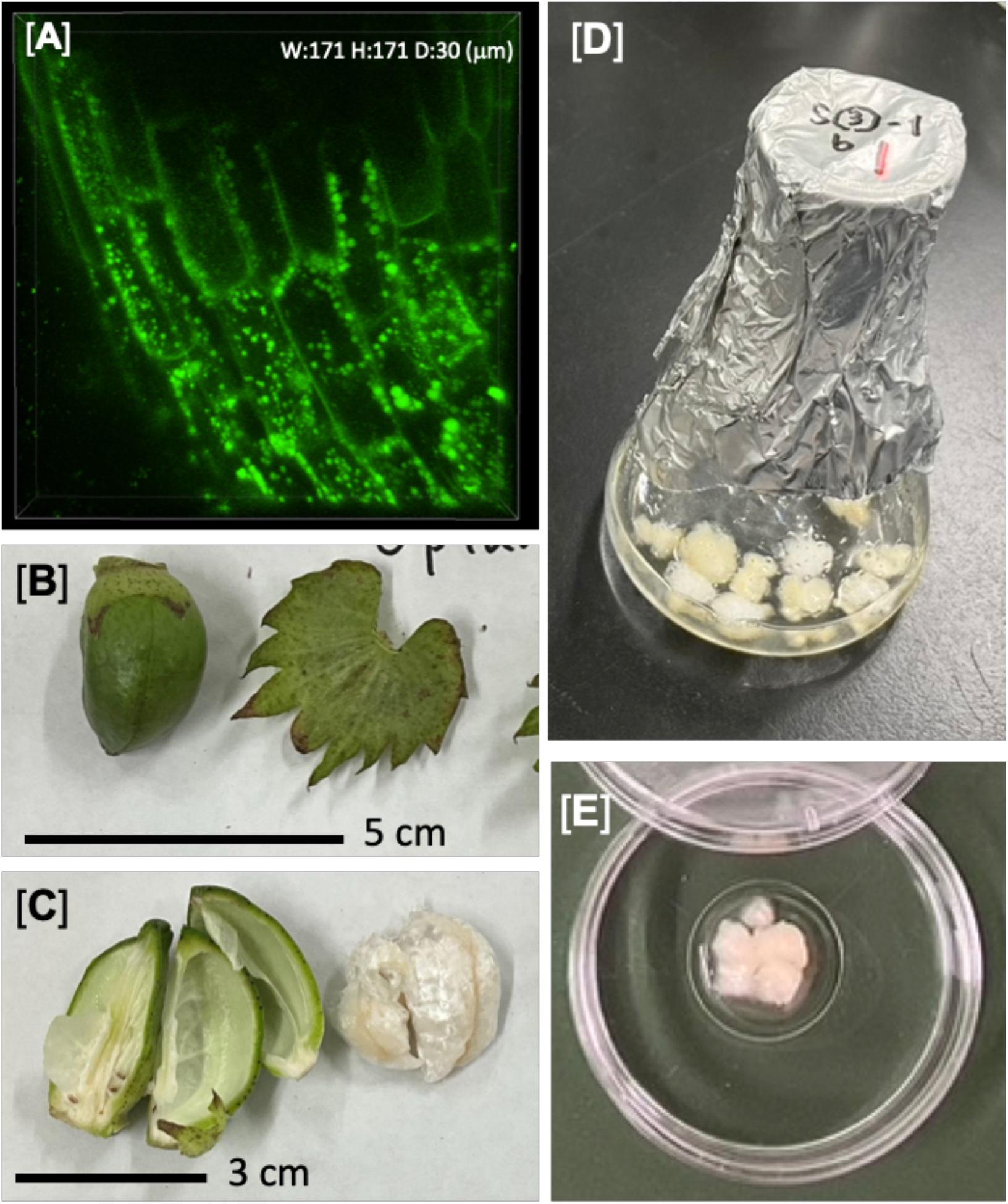
Plant materials for the experiments. [**A**]; Fluorescent substances appeared on the root epidermal cells of *Arabidopsis thaliana* with the probe treatment (2 x 10^−5^ M, 30 min). The 3-D view image was built from 31 slice views with maximum intensity projection. Z-axis stack = 1 μm/step). Size of the 3-D view is shown at the top. [**B]**; Cotton boll of *G. hirsutum* (30 DPA). [**C**]; Cotton fibers (right) in the boll (*G. hirsutum*) in [B]. [**D**]; Cotton ovule culture (*G. arboreum*, cultured for 10 days) prepared in a 50 ml Erlenmeyer flask. [**E**]; Cotton cultured ovules placed in a bottom flat petri dish.

### 2. Preparation of probe solution and application

The probe molecule was synthesized as described (Tamogami et al., 2021). Probe molecule (4.6 mg) was dissolved in DMSO (5 ml) to give a probe solution of 2 x 10^−3^ M. Diluted probe solution (2 x 10^−5^ M) with water was prepared, and cotton fibers/ovules were dipped in this solution. After an appropriate incubation time period at 25ºC, the plant materials were rinsed with distilled water (3 times) and transferred to a bottom flat petri dish (1 cm diameter, IWAKI Glass Company Ltd., Japan). The petri dish was set on the microscopy table and observed (see below 3.). Control plant materials were treated with the test solution without the probe. In case of *Arabidopsis thaliana* (Columbia), seeds were placed in a glass dish (6 cm diameter) with water (2 ml) and incubated for 3 days (25ºC, light/dark cycle = 12/12 hrs). *Arabidopsis* seedlings were then transferred to a bottom flat petri dish. And 0.25 ml of the probe solution (2 x 10^−5^ M) was applied. Plant materials were fully dipped in the solution. After incubation for 30 min, plant materials were rinsed with water (3 times) and observed under a microscope (see below 3.).

### 3. Equipment and creating 3-D view images of the targets

Fluorescence microscopy images were obtained using a Nikon Confocal Fluorescence Microscope System AX (Nikon Corporation, Japan). Excitation wavelength = 488 nm. Emission wavelength = 499 - 551 nm. Transmitted-light view was obtained at 488 nm. Using ND acquire software (supported by Nikon Corporation) to obtain 3-D view images, plant materials were scanned for an adequate period of time. Three-D view-images were prepared with maximum intensity projection by the software (NIS-Elements imaging software) provided by the manufacturer (Nikon Corporation, Japan).

## Results and discussion

### 1. View images of fluorescent substances in the cotton fibers

The development of cotton fibers can be roughly divided into three stages: fiber initiation, elongation and secondary cell wall biosynthesis, and the biosynthesis been proposed to be active for around 30 DPA (Lee et al., 2007). We used cotton fibers of *G. hirsutum* (upland or American cotton) in the bolls on 30 DPA for the experiments. Cotton fibers were treated with the probe, and observed under a fluorescence microscope. Figure **2** [**A**] showed a typical 3-D view image obtained from the fibers of *G. hirsutum* treated with the probe. Three-dimensional images were useful for observing the overall progression of how these fluorescent substances were localized. The pericellular lines (possibly, the primary cell walls) of elongated fibers were recognized as the fluorescent lines composed of small particles. It should be noted that the fluorescent substances were clearly found to be localized in the cotton fibers. From the typical slice view (Fig. **2** [**B**]), it was clear that the fibers of *G. hirsutum* included many fluorescent substances as characteristic fragments or zones.

**Fig. 2.**
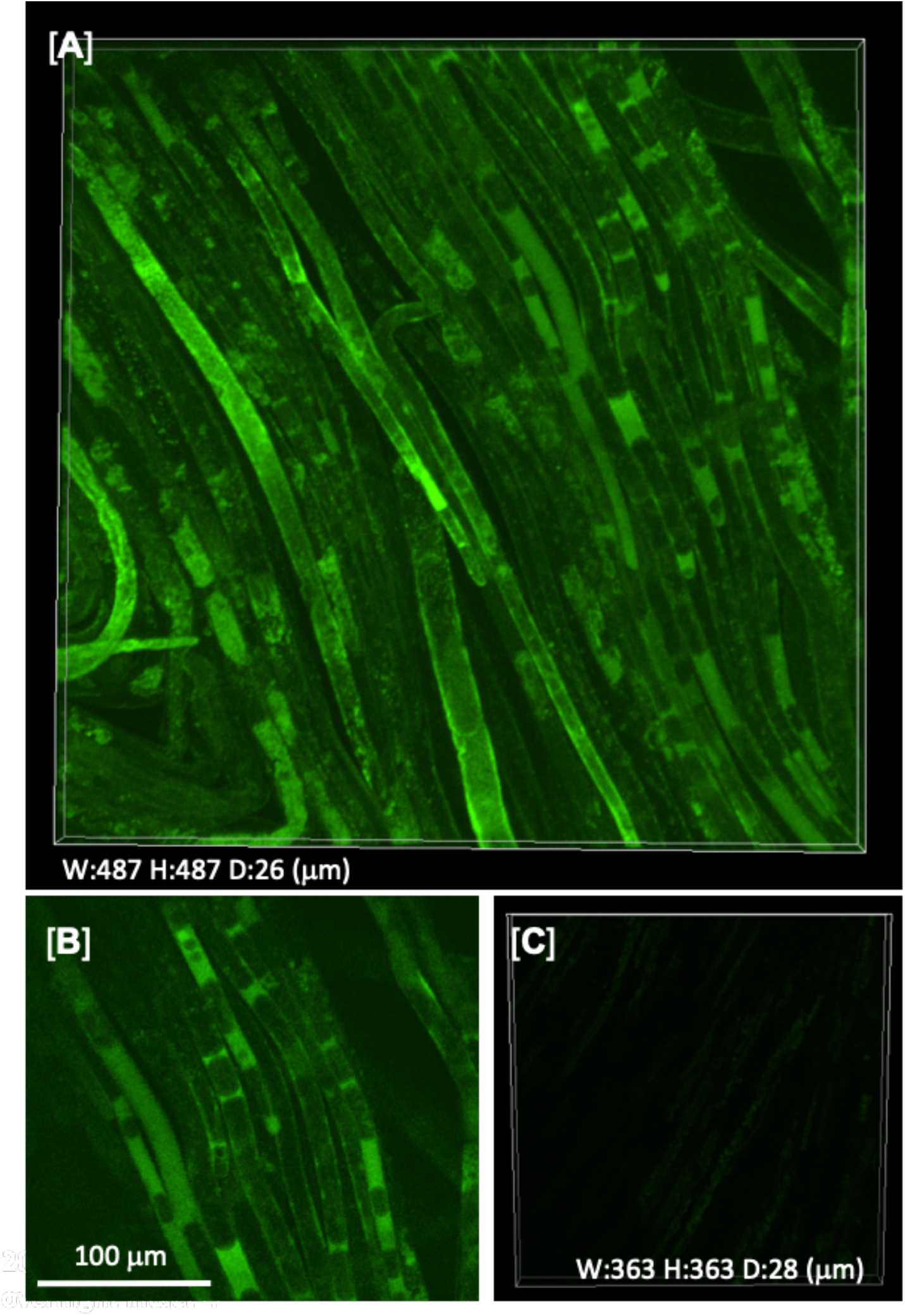
Fluorescent substances in the cotton fibers of *G. hirsutum* (30 DPA) treated with the probe. [**A**]; 3-D view image of the fluorescent products in cotton fibers treated with the probe (2 x 10^−5^ M, overnight). Fluorescent substances were observed as fluorescent straw tubes. The view was constructed from 27 slice views with maximum intensity projection. Z-axis stack = 1 μm/step. Size of the 3-D view (x, y, z-axis) is shown at the bottom left. [**B**]; Representative slice view of the cotton fibers in [**A**]. [**C**]; View of the cotton fibers treated with the test solution without the probe.

We then tested the probe on fibers of *G. arboreum* (Asian cotton) in the bolls on 25 DPA (Fig. **3**, [**A**] and [**B**]). The pericellular lines (possibly, the primary cell walls) of the fibers treated with the probe were clearly observed, and they appeared to be composed from the small fluorescent particles like in *G. hirsutum*. However, fluorescent substances in these fibers appeared to be packaged, or formed intricate spirals (Fig. **3**, [**A**] and [**B**]). It was interesting that these two varieties were different from each other in shapes and localization manners with the fluorescent substances in their fibers. Fluorescent bands and zones might be characteristics of the *G. hirsutum* variety. The presence of plentiful fluorescent products in the cotton fibers suggested to us the usability of this probe. In addition, this probe might be used as a reagent to characterize the quality of the cotton fibers.

**Fig. 3.**
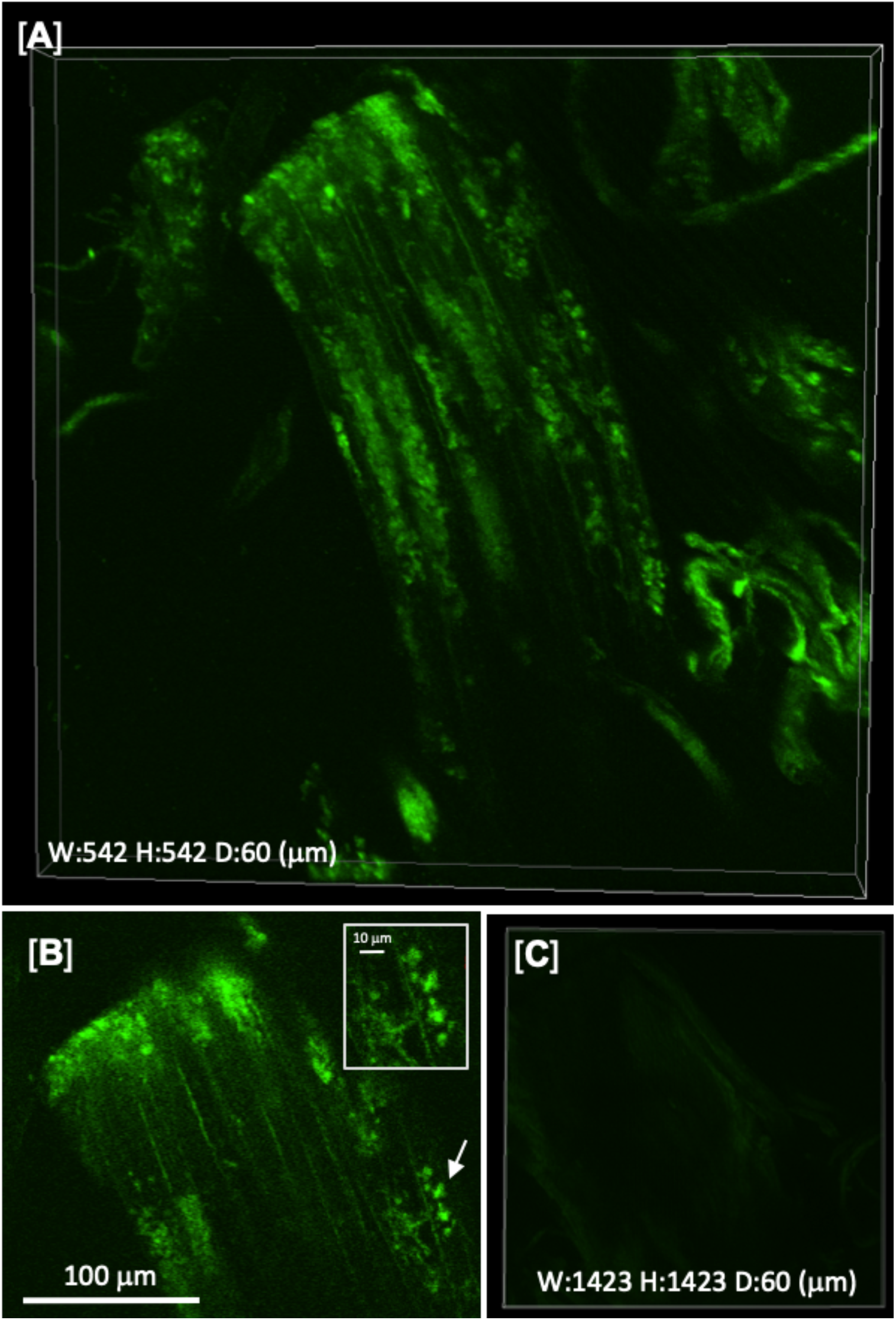
Fluorescent substances in the cotton fibers of *G. arboreum* (25 DPA) treated with the probe. [**A**]; 3-D view image of the fluorescent products in the cotton fibers treated with the probe (2 x 10^−5^ M, overnight). The cotton fibers were observed like a fluorescent waterfall. The view was constructed from 61 slice views with maximum intensity projection. Z-axis stack = 1 μm/step. Size of the 3-D view (x, y, z-axis) is shown at the bottom left. [**B**]; Representative slice view of the cotton fibers in [**A**] with a magnified view of the fluorescent substances at right below (an arrow). [**C**]; View of the cotton fibers treated with the test solution without the probe.

### 2. View images of fluorescent substances in the cultured ovules

We prepared cotton ovule cultures according to a previous method (Tan et al., 2013), and observed ovules treated with the probe. On the surface of ovules (*G. hirsutum*, 3 days culture) treated with the probe, many circular fluorescent substances were observed (Fig. **4** [**A**]). Most of these substances were double-ring structures, and some small single-ring substances were also identified. It was found that fluorescent particles are localized inside of the rings, and some of them appeared to be emitting from small holes in the center (such as, **a** and **b**). Skeletons of the ring-like substance (**f** in [**D**]) could only be slightly recognized in the transmitted-light view [**E**], and the ring-like substance existed on the cracks on the ovule in the merged view [**F**]. It appeared that these ring-like substances came from the cracks on the ovule surface. These ring-like substances appeared similar to the cotton stomata on the ovules (Stewart, 1975), but it was not clear whether these substances were the stomata, or not.

**Fig. 4.**
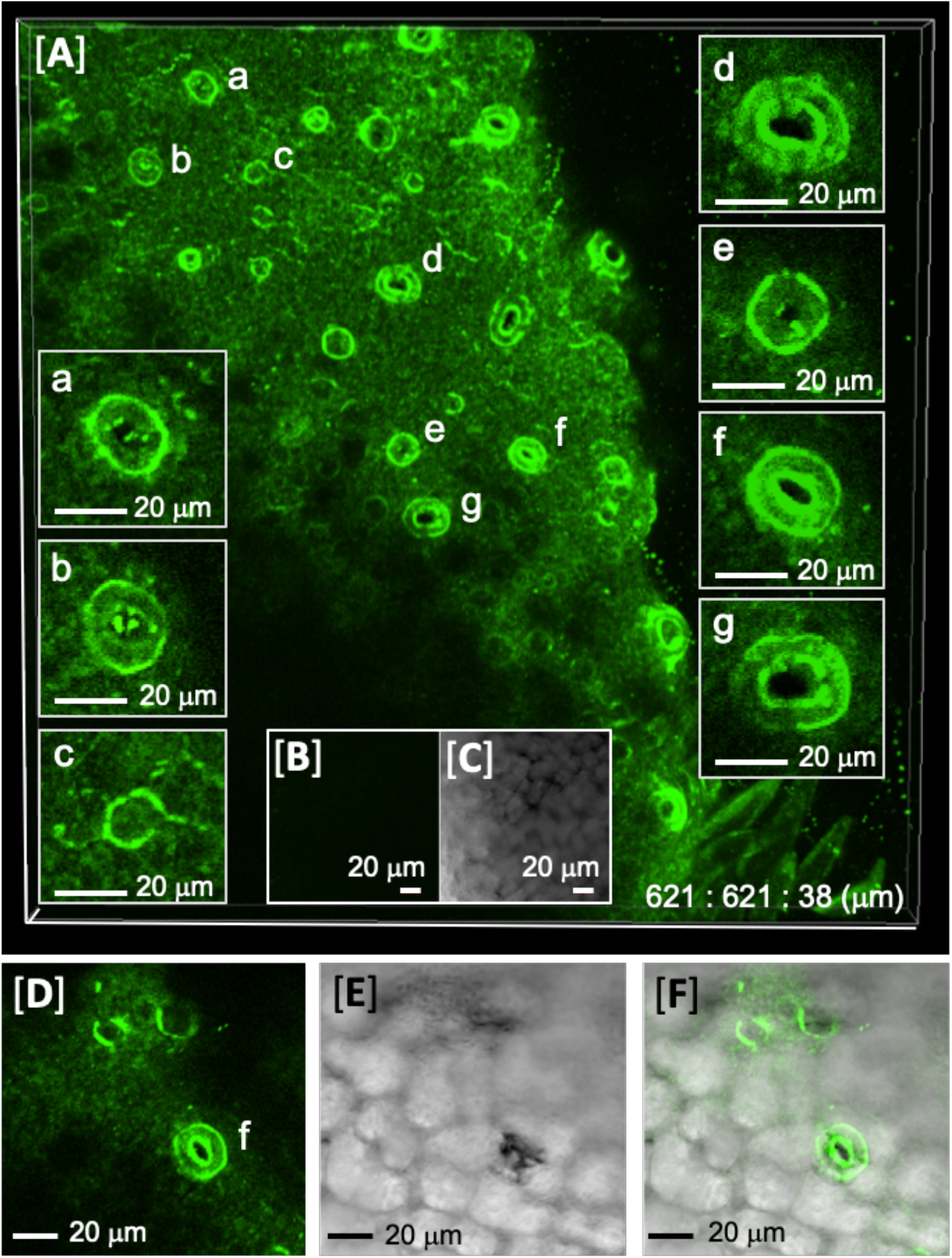
Fluorescent ring-like structures on the cotton cultured ovule of *G. hirsutum* (3 days culture) treated with the probe (2 x 10^−5^ M, 3 hrs). [**A**]; 3-D view image constructed from 39 slice views with maximum intensity projection. Z-axis stack = 1 μm/step. Size of the 3-D view (x, y, z-axis) is shown at the bottom right. Typical slice views (**a** - **g**) were shown in the squares. [**B**]; Control view of *G. hirsutum* (3 days culture) treated with the test solution without the probe. [**C**]; Control transmitted-light view. Scale bar = 20 μm. [**D**]; Slice view of fluorescent ring-like structures (**f**) in [**A**] from another view-angle. [**E**]; Transmitted-light view. [**F**]; Merged view.

Conversely, on the cultured ovule of *G. arboreum* (10 days culture), short but strong looking fibers were found as shown in Fig. **5** [**A**]. Presumably, this area is near to the end of the ovule, because of less fiber initiation (Stewart, 1975). Spiral patterns were recognized at the top of the fiber (marked with **a**) as shown in the enlarged view (Fig. **5** [**B**]). In other fibers, fluorescent particles of band-like shapes (**b**) were recognized as shown within a square in Fig. **5** [**A**]. These spirals might be typical with early short fibers of *G. arboreum*. It was clear that the fiber on the left in Fig. **5** [**A**] originated from a polygonal hole (marked with an asterisk), which appeared to be slightly raised with surrounding cells (Fig. **5** [**C**], [**D**]). Moreover, a very characteristic fluorescent structure was observed on another ovule treated with the probe (Fig. **5** [**E**] - [**I**]). This polygonal structure was surrounded by eight epidermal cells to be an octagon, and its skeleton and contents were very clear. From the 3-D view image (Fig. **5** [**E**]), it was found that the spirals (indicated by an arrow) were included in this octagon. Slice views showing the upper part [**F**], lower part [**G**], and middle part [**H**] of the octagon, suggested that the spirals might be accumulated in the middle part [**H**]. In the transmitted-light view [**I**], only a faint outline of the octagonal frame was recognized. Possibly, the spirals might be embedded beneath the surface. It is likely that plant hormones added to the culture, as well as aqueous growth condition, might cause morphological changes in ovule development, and the cultured ovule cells grew differently from natural ovules. Detailed studies on these unique substances (Fig. **4** and **5**) are currently under investigation.

**Fig. 5.**
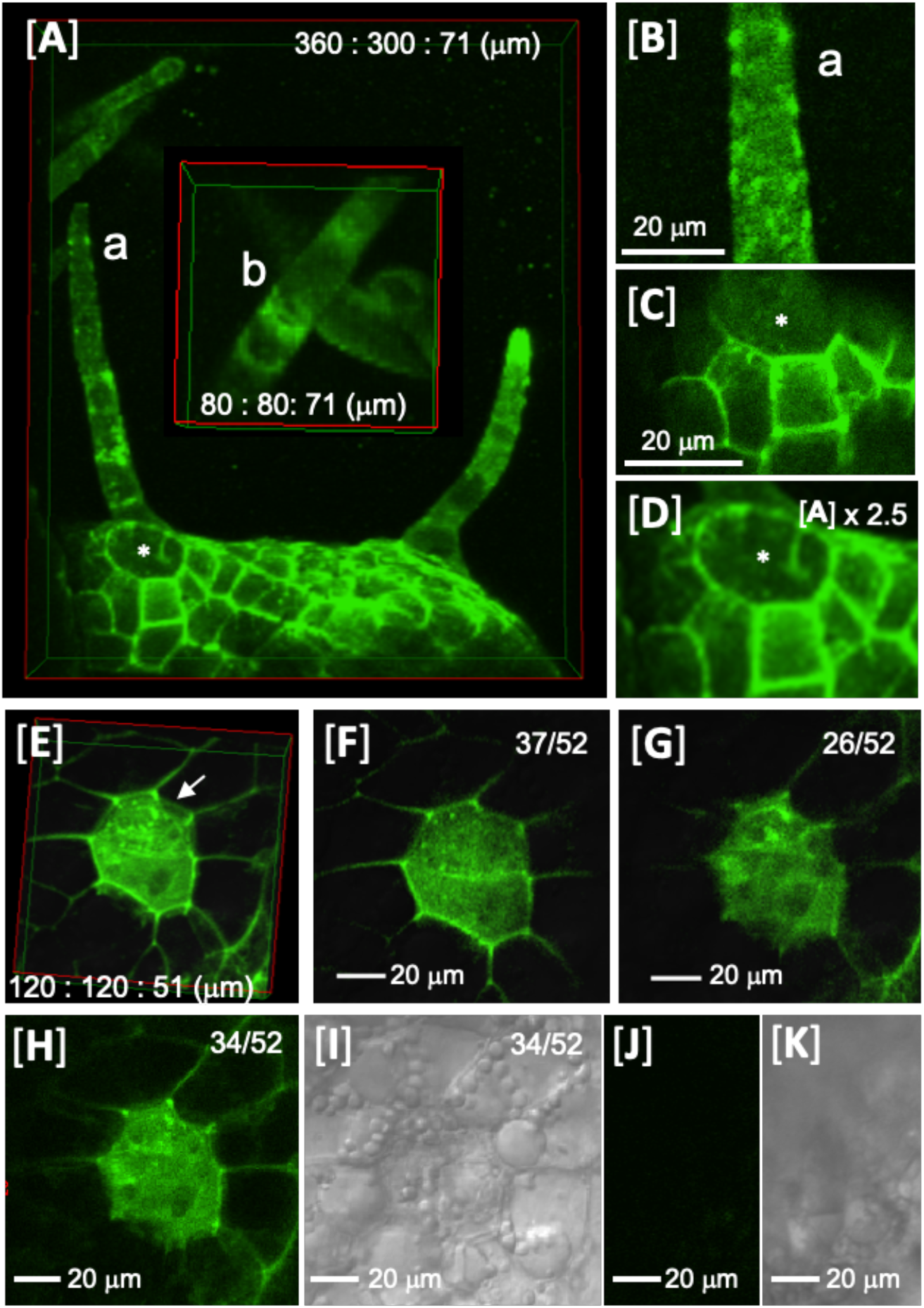
Fluorescent substances appeared on the cotton cultured ovules of *G. arboreum* (10 days culture) treated overnight with the probe ([**A**] - [**D**]), or 30 min ([**E**] - [**I**]). [**A**]; Fluorescent substances on the cotton cultured ovules treated with the probe (2 x 10^−5^ M, overnight). The view was constructed from 72 slice views with maximum intensity projection. Z-axis stack = 1 μm/step. Size of the 3-D view (x, y, z-axis) is shown at the upper right. In square **b**, typical fluorescent stripes in the fiber were shown. [**B**]; Spirals in the fiber (**a**) in [**A**]. [**C**]; Representative slice view of the polygonal hole (marked with an asterisk). [**D**]; Enlarged view of the polygonal hole. Views of [**E**] - [**H**] showed fluorescent octagon on the cotton cultured ovule treated with the probe. [**E**]; 3-D view image built from 52 slice views with maximum intensity projection. Z-axis stack = 1 μm/step. Size of the view (x, y, z-axis) is shown at the bottom. [**F**]; Upper slice view of [**E**] at 37th from the top. [**G**]; Lower slice view of [**E**] at 26th from the top. [**H**]; Middle slice view of [**E**] at 34th from the top. [**I**]; Transmitted-light view of [**H**]. [**J**]; Control fluorescence view (without the probe). [**K**]; Transmitted-light view of ([**J**]).

In summary, the ability of the labeling probe was examined with cotton fibers and cultured ovules. Various fluorescent substances were clearly found in/on them. Elucidation of these fluorescent substances, and investigations if they actually engage in cellulose biosynthesis are now under way.

## Acknowledgements

We appreciate Dr. M. Abe (Akita Prefectural University) for growing plant materials of *G. arboreum*, and Dr. K. Sakurai (Akita Prefectural University) for his kind gift of *G. hirsutum* flowers and bolls. We appreciate Akita Prefectural University for the use of fluorescence microscope equipment to observe the samples, and analytical equipment, such as NMR and MS for the probe synthesis.

## Author contribution

S. T. prepared the probe compound, designed experiments, carried out microscopic experiments, wrote a draft and prepared the manuscript; A. W. designed experiments, prepared cotton ovule cultures, analyzed experimental results and discussed with S, T; G. K. A. and R. R. discussed results with S. T. and edited the manuscript draft.

